# EnClaSC: A novel ensemble approach for accurate and robust cell-type classification of single-cell transcriptomes

**DOI:** 10.1101/754085

**Authors:** Xiaoyang Chen, Shengquan Chen, Rui Jiang

## Abstract

**Background:** In recent years, the rapid development of single-cell RNA-sequencing (scRNA-seq) techniques enables the quantitative characterization of cell types at a single-cell resolution. With the explosive growth of the number of cells profiled in individual scRNA-seq experiments, there is a demand for novel computational methods for classifying newly-generated scRNA-seq data onto annotated labels. Although several methods have recently been proposed for the cell-type classification of single-cell transcriptomic data, such limitations as inadequate accuracy, inferior robustness, and low stability greatly limit their wide applications.

**Results:** We propose a novel **en**semble approach, named EnClaSC, for accurate and robust cell-type **cla**ssification of **s**ingle-**c**ell transcriptomic data. Through comprehensive validation experiments, we demonstrate that EnClaSC can not only be applied to the self-projection within a specific dataset and the cell-type classification across different datasets, but also scale up well to various data dimensionality and different data sparsity. We further illustrate the ability of EnClaSC to effectively make cross-species classification, which may shed light on the studies in correlation of different species. EnClaSC is freely available at https://github.com/xy-chen16/EnClaSC.

**Conclusions:** EnClaSC enables highly accurate and robust cell-type classification of single-cell transcriptomic data via an ensemble learning method. We expect to see wide applications of our method to not only transcriptome studies, but also the classification of more general data.

## Background

Recent advances in single-cell RNA-sequencing (scRNA-seq) techniques make it possible to reveal previously unknown heterogeneity and functional diversity at a microscopic resolution [1-3]. The exponential growth of the number of cells profiled in individual scRNA-seq experiments has shed light on the studies aiming to identify new cell types [4, 5], reveal regulatory mechanisms [6, 7], assess tissue composition [1, 8, 9], investigate cell development and lineage processes [10-12], and many others. Currently, most of the analysis methods for scRNA-seq data are commenced with unsupervised clustering, which highly relies on the investigator’s background knowledge about the signature molecules, and is not efficient and accurate enough for the cell-type assignment of clusters [13]. Therefore, there is a demand for novel computational methods for classifying newly-generated scRNA-seq data onto annotated labels.

A variety of methods have recently been proposed for the cell-type classification of single-cell transcriptomes. For example, an unsupervised approach, named scmap [14], projects cells to the identified cell-types based on the similarity between query and reference cells. Conventional supervised learning-based methods, such as Random Forest and Support Vector Machine, though can be used for the cell-type classification, have been proven that their performance is not comparable to scmap. SuperCT, a supervised neural network framework, characterizes cell types of single-cell transcriptomic profiles with transformed binary features. Nevertheless, there are still several limitations to be addressed in the proposed methods for the cell-type classification of single-cell transcriptomes. First, even the state-of-the-art methods have achieved encouraging performance, the classification performance and stability can still be further improved for various datasets and tasks as shown in the Results Section. Second, to select informative features of the query and reference datasets, there is a demand for a tailored feature selection approach for the cell-type classification. Third, both the query and reference datasets may contain cell types that have only a small number of samples. A superior classification method should be able to effectively characterize the cell types with only a small number of samples. Last but equally important, a method that can be generally applied to data of various dimensions or dropout rates is desirable for the broader application scenarios.

Motivated by the above understanding, we propose in this paper EnClaSC, a novel **en**semble approach for accurate and robust cell-type **cla**ssification of **s**ingle-**c**ell transcriptomes. Through comprehensive experiments, we illustrate that our method is superior to existing methods in not only the self-projection within a specific dataset, but also the cell-type classification across datasets. With a few-sample learning strategy, EnClaSC can effectively characterize the cell types that have only a small number of samples. We further show the robustness of our method for various data dimensionality and different data sparsity. Through joint analysis of classification results with scRNA-seq datasets of different species, we demonstrate the ability of our method to make the cross-species cell-type classification.

## Methods

### Design of EnClaSC

As illustrated in Fig. 1, EnClaSC consists of four modules. First, a feature selection module finds informative genes which benefit the cell-type classification from the common genes of the query and reference sets. Second, a few-sample learning module is adopted to sufficiently learn the characteristics of classes with few samples, and thus improve the performance in identifying the few-sample classes. Third, a neural network module uses artificial neural networks with an ensemble learning strategy to stabilize the performance of the cell-type classification. Finally, a joint prediction module integrates outputs of the few-sample learning and neural network modules to predict the class that a query cell belongs to. In general, EnClaSC draws on the idea of ensemble learning in the feature selection, few-sample learning, neural network and joint prediction modules, respectively, and thus constitutes a novel ensemble approach for cell-type classification of single-cell transcriptomes.

**Fig. 1.**
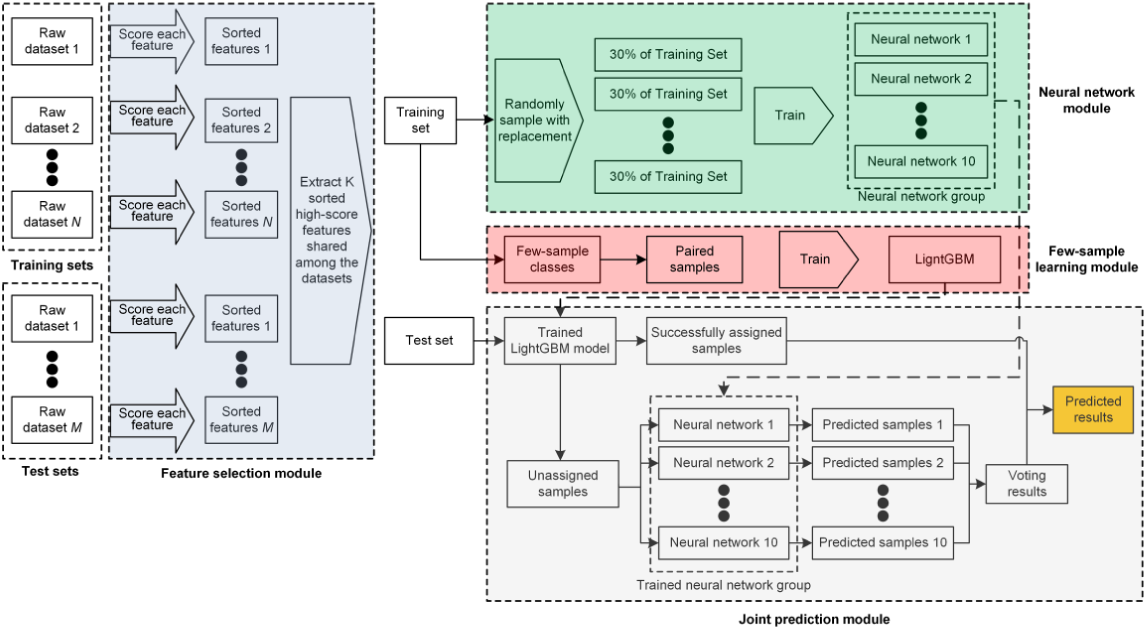
The graphical illustration of EnClaSC. EnClaSC consists of four modules, including feature selection, few-sample learning, neural network and joint prediction modules.

### Feature selection

Let *E* be the value of gene expression in scRNA-seq, *n* the number of the cells in the dataset. The dropout rate of the *j th* feature is abbreviated as *D*_*j*_. For the *i th* cell (sample), the expression of the *j th* feature (gene) of is abbreviated as *E*_*ij*_. For the *j th* feature, the level of gene expression *F*(*j*) is calculated by the logarithm-plus-one of its arithmetic mean value as

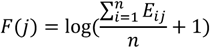

For the estimation of gene expression, we first use the dropout rate to fit a linear model using the least square method

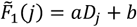

We then estimate *F*(*j*) by the mean of the logarithm-plus-one of each feature, namely,

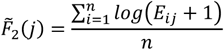

The residuals of these two approaches between the true gene expression level are recorded as Δ*F*_1_(*j*) and Δ*F*_2_(*j*), respectively. Briefly, 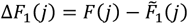, and 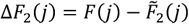.ΔF_1_(*j*) reflects the rate of gene expression, that is, whether the gene can be expressed or not. Δ*F*_2_(*j*) is the information entropy of each gene, which reflects the residual degree of each gene expression. We can use the integrated score *G*(*j*) = *α*Δ*F*_1_(*j*) + (1 – *α*)Δ*F*_2_(*j*) to consider the above two factors synthetically. Here, *α* is the control coefficient, which is responsible for regulating the influence of the two calculated factors. We set *α* to 0.5 in all the experiments of this work.

For a specific scRNA-seq dataset, we compute *G*(*j*) as the scores of features, and then sort the features according to the scores in descending order. We extract *K* sorted high-score features shared among the training and test sets. This feature selection approach can effectively maximize the common high-score features of different scRNA-seq datasets, and thus contribute to the cell-type classification across datasets.

### Few-sample learning strategy

We define few-sample classes as the cell types whose number of samples does not exceed 0.5% of the total number of training samples. In order to fully extract features of the few-sample classes, we perform data augmentation and pre-train a few-sample classification model using samples of the few-sample classes in the training set. In more detailed, let *N* be the number of samples in the training set, n the number of samples of the few-sample classes in the training set, *M* the number of samples in the test set, and *D* the number of features after feature selection. We pair the samples of these few-sample classes one by one to form a few-sample training set with *n* × *n* samples and 2 × *D* features. If the two cells of a paired sample belong to the same cell type, we mark the label of the paired sample as 1; otherwise, we mark it as 0. We then pair the samples of the few-sample classes in the original training set with each sample in the original test set to form a few-sample test set with *n* × *M* samples and 2 × *D* features.

We adopt LightGBM, a gradient boosting framework that uses tree-based learning algorithms, to perform the few-sample training. We use the default setting of parameters except for the parameters listed in Table 1 [15]. To obtain the probability scores that an original test sample belongs to the few-sample classes, we weight the prediction results of the few-sample test set by the *Pearson* correlation coefficients between the expression values of the two cells in corresponding few-sample test samples drawing on the idea of ensemble learning, namely,

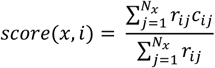

where *x* is one of the classes in the original training set, *N_x_* the number of samples of class *x* in the original training set, *j* the *j th* sample of class *x* in the original training set, *i* the *i th* sample of class *x* in the original test set, and *r*_*ij*_ the *Pearson* correlation coefficients between *i* and *j, c*_*ij*_ the prediction result of LightGBM for the paired sample of *i* and *j*.

**Table 1.** Parameters of the LightGBM model.

| Parameters | Setting |
| --- | --- |
| boosting_type | 'gbdt' |
| objective | 'regression' |
| metric | {'l2', 'auc'} |
| learning_rate | 0.05 |
| bagging_freq | 5 |
| verbose | 1 |

**Table 2.** The architecture of the neural network.

| Layers | Setting |
| --- | --- |
| Dense_1 | {(feature numbers,128), 'relu'} |
| Dropout_1 | 0.25 |
| Dense_2 | {(128,64), 'relu'} |
| Dropout_2 | 0.5 |
| Dense_3 | {(64,32), 'relu'} |
| Dense_4 | {(32,number of classes in training sets), 'softmax'} |

If the predicted maximum score of an original test sample is greater than *γ*, then we assign the sample as the class that has the maximum score, otherwise, the sample is predicted as “unassigned” and should be further classified using the subsequent artificial neural networks.

### Artificial neural networks for the cell-type classification

The neural network module uses artificial neural networks with ensemble learning strategy to classify the test samples which are predicted as “unassigned” by the few-sample learning module. We first design an artificial neural network framework with parameters as shown in Table 1, and implement it using Keras with Tensorflow as the backend. In order to improve the stability of the classification performance, drawing on the Bootstrap strategy, we generate 10 new training sets by selecting 30% of the original training set at random with replacement, and then use them to train 10 neural networks, respectively. For each trained neural network, if the predicted maximum score of a test sample exceeds, again, *γ*, the sample is assigned as the class that has the maximum score, otherwise, it is classified as “unassigned”. We set *γ* to 0.7 in all the experiments of this work. For each test sample, we classify it as “unassigned” unless there are more than half of the 10 neural networks predict the test sample as a specific same class, and we thus classify the sample based on the voting result of the 10 neural networks.

### Assessment of performance

We adopt the widely used *kappa* value and *assigned rate* value to assess the classification performance of the models. Briefly, we calculate the *kappa* values of the classification results of assigned samples and the real classes of corresponding samples using the following formula

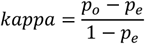

where *p*_0_ is the sum of the number of samples correctly classified divided by the number of samples in the test set, that is, the overall classification accuracy of the test set. Assuming that the number of real samples in each class of test set is *a*_1_, *a*_2_ … *a*_*C*_ respectively, the number of samples in each class of the test set predicted is *b*_1_, *b*_2_ … *b*_*C*_ respectively, and the number of samples in the test set is *n*. We have

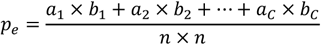

For the assessment of the ability of a model to recognize the cells in the test set, we use *assigned rate* to represent the proportion of the number of samples that can be assigned to a cell type by the classifier, namely,

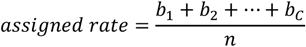

Because the classes of test sets and training sets may not be completely consistent, the *assigned rate* should be maintained at a relatively high level, but not the higher the better, and thus the classification performance is primarily related to *kappa* values.

### Data collection

We downloaded three humans pancreatic scRNA-seq dataset provided by Baron, M. et al., Muraro, MJ et al., and Xin, Y. et al. (hereinafter are abbreviated as Baron Dataset, Muraro Dataset, and Xin Dataset, respectively) from NCBI Gene Expression Omnibus via accession GSE84133, GSE85241, and GSE81608, respectively [16-18]. The human pancreatic scRNA-seq dataset provided by Segerstolpe, Å. et al. (hereinafter is abbreviated as Segerstolpe Dataset) was downloaded from EMBL-EBI ArrayExpress via accession E-MTAB-5061 [5]. For the subsequent usability of the dataset, we removed the cells of the “unclear” class in Muraro Dataset, the cells of the “not applicable”, “unclassified endocrine” and “unclassified” classes in Segerstolpe Dataset and the cells of the “alpha.contaminated”, “beta.contaminated”, “delta.contaminated”, and “gamma.contaminated” classes. After preprocessing, Baron Dataset contains 8569 samples with 20, 125 features, Muraro Dataset contains 2122 samples with 19, 127 features, Xin Dataset contains 1492 samples with 39, 851 features, and Segerstolpe Dataset contains 2166 samples with 25, 525 features.

We also downloaded two mouse retina scRNA-seq datasets provided by Macosko, EZ et al. and Shekhar, K. et al., (hereinafter are abbreviated as Macosko Dataset and Shekhar Dataset, respectively) from NCBI Gene Expression Omnibus via accession GSE63473 and GSE81904 [2, 19]. Macosko Dataset measures the expression of 23, 288 genes in 44, 808 cells, while Shekhar Dataset measures the expression of 13, 166 genes in 27, 499 cells.

Two mouse brain cell scRNA-seq datasets provided by Romanov, RA et al. and Zeisel, A. et al. (hereinafter are abbreviated as Romanov Dataset and Zeisel Dataset, respectively) and one mouse brain cell scRNA-seq dataset collected by Darmanis, S. et al. (hereinafter is abbreviated as Darmanis Dataset) were downloaded from the NCBI Gene Expression Omnibus via accession GSE74672, GSE60361 and GSE67835, respectively [9, 20, 21]. To unify the label information, we replaced the cell type label ‘oligos’ in the Remanov Dataset with ‘oligodendrocytes’, and the cell type labels ‘ca1pyramidal’, ‘s1pyramidal’ and ‘interneurons’ in the Zeisel Dataset with ‘neurons’. After preprocessing, Romanov Dataset contains 2881 samples with 24, 341 features, Zeisel Dataset contains 3005 samples with 19, 972 features, and Darmanis Dataset contains 466 samples with 22, 088 features.

## Results

### EnClaSC achieves high performance for self-projection within a dataset

In order to illustrate that our method can effectively perform self-projection within a scRNA-seq dataset, we conducted a series of self-projection experiments using 6 datasets, including Baron Dataset, Muraro Dataset, Xin Dataset, Segerstolpe Dataset, Macosko Dataset and Shekhar Dataset. We compared the performance of our method with scmap, a similarity-based method, and SuperCT, an ANN-based method (we self-implemented the method according to the paper because SuperCT is not open source) [13, 14]. Using the same training (reference) and test (query) sets with EnClaSC, we run the two baseline methods with parameters or structures proposed by the respective authors.

Both EnClaSC and scmap provide a feature selection method, while SuperCT uses all features. When running EnClaSC and scmap, we selected 100 features, which is considered to be able to better demonstrate the advantages of scmap in their paper. We performed 5-fold cross-validation within each of the six datasets. EnClaSC model has better self-projection performance than SuperCT and scmap. At the same time, in terms of self-projection stability, EnClaSC is significantly superior to SuperCT. The variance of the *kappa* value of EnClaSC is 95.05% lower than SuperCT, and the variance of the *assigned rate* of EnClaSC is 85.60% lower than SuperCT.

### EnClaSC outperforms other methods in the cell-type classification across datasets

We use six scRNA-seq datasets to demonstrate the superior cell-type classification performance of EnClaSC. Using Baron Dataset, Muraro Dataset, Xin Dataset, and Segerstolpe Dataset these four human pancreatic datasets, we selected each one of them as the test set, while the integration of remaining three datasets serves as the training set, respectively. In addition, we used two mouse retina datasets, namely, Macosko Dataset and Shekhar Dataset, to form two symmetric training-test sets. We used 100 selected features in EnClaSC and scmap, while all features in SuperCT, and repeated each experiment five times.

As shown in Fig. 3, the *kappa* value of EnClaSC is higher than that of scmap given comparable *assigned rates*, except for the group where Macosko Dataset serve as the test set and scmap achieves much lower *assigned rates*. Compared with SuperCT, which also uses a neural network architecture, *assigned rates* of EnClaSC has little difference except for Baron Dataset as the test set, while most *kappa* values of EnClaSC outperform SuperCT. When Baron Dataset serves as the test set, the *assigned rates* of SuperCT is much lower than that of scmap and EnClaSC. In addition, it can be seen from the figure that the classification performance of EnClaSC is much more stable than that of SuperCT. Therefore, even though EnClaSC only uses 100 features, the classification performance of EnClaSC is still far better than that of SuperCT. In general, EnClaSC has a more outstanding performance in cell-type classification compared with other two base-line methods.

**Fig. 2.**
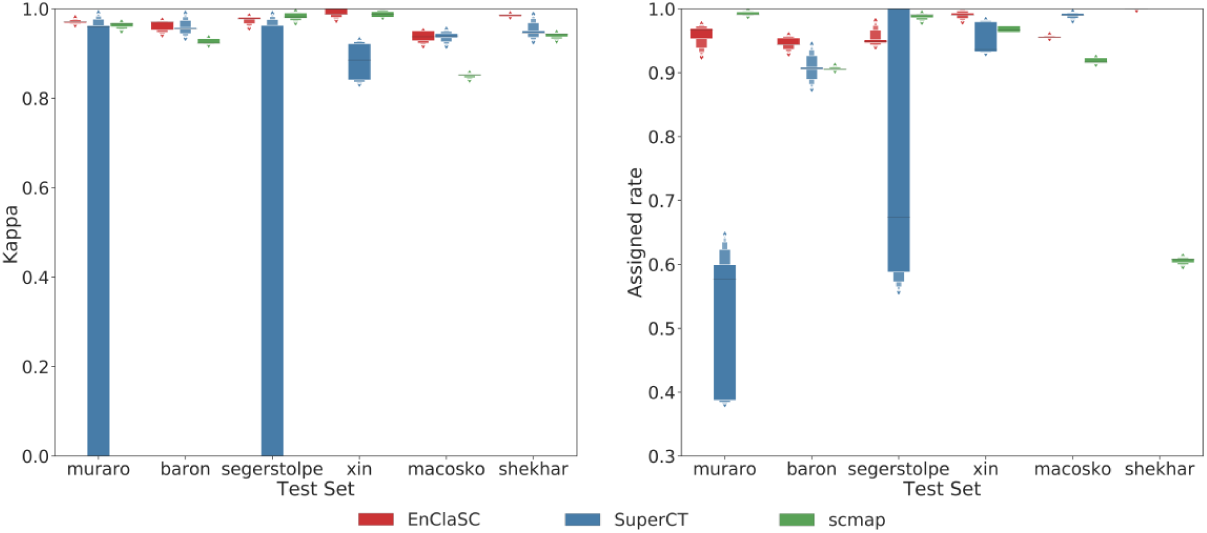
Performance comparison of the self-projection within a specific dataset.

**Fig. 3.**
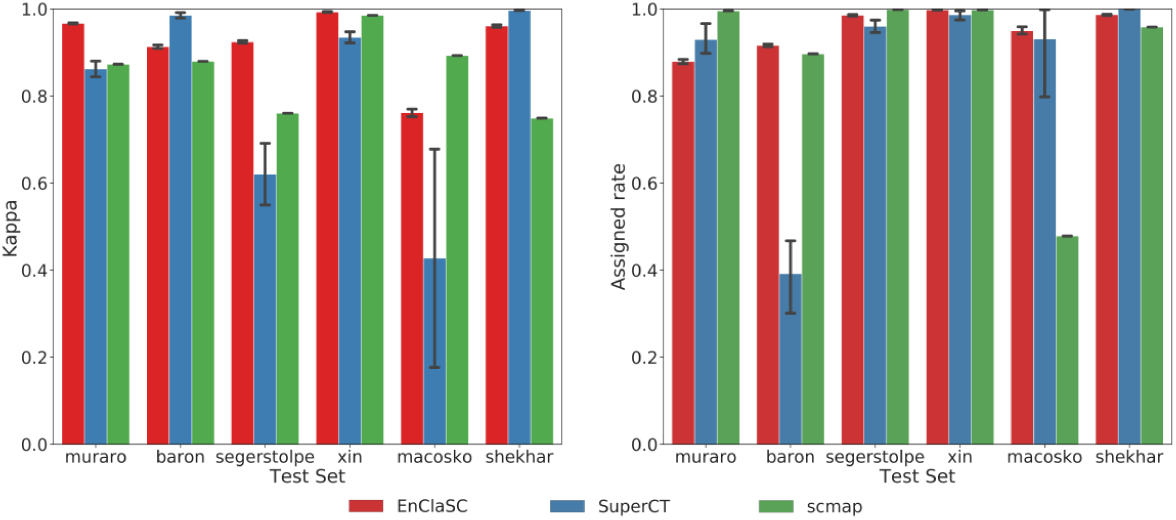
Performance comparison of the cell-type classification across different datasets.

### Contribution of each module

#### Feature selection module

We validated the contribution of the feature selection module using the four human pancreatic datasets including Baron Dataset, Muraro Dataset, Xin Dataset and Segerstolpe Dataset, and two mouse retina datasets including Macosko Dataset and Shekhar Dataset. We performed six experiments as the above section. We first took each one of the four human pancreas datasets as a test set and integrated the remaining three datasets as a training set. Then we took each one of the two mouse retina datasets as a test set, and the remaining one was served as a training set. We replaced the feature selection method of our feature selection module with PCA, which is commonly used for scRNA-seq dimensionality reduction, feature selection methods of Seurat v3.0 and that of scmap, and compared them with our original method [22].

The results are shown in Fig. 4. Compared to PCA, our feature selection method is more stable. The *kappa* and the *assigned rate* of EnClaSC is much better than the EnClaSC framework using the feature selection method of Seurat v3.0 (the average of *kappa* values of EnClaSC is 69.65% higher than Seurat v3.0, and the average of *assigned rates* is 3.95% higher than Seurat v3.0). EnClaSC has slightly higher *kappa* values with comparable *assigned rates* compared with the EnClaSC framework using the feature selection method of scmap. In summary, the feature selection module of EnClaSC is superior to other widely used feature selection approaches in the EnClaSC framework, and thus serves as the effective feature selection approach for the cell-type classification.

**Fig. 4.**
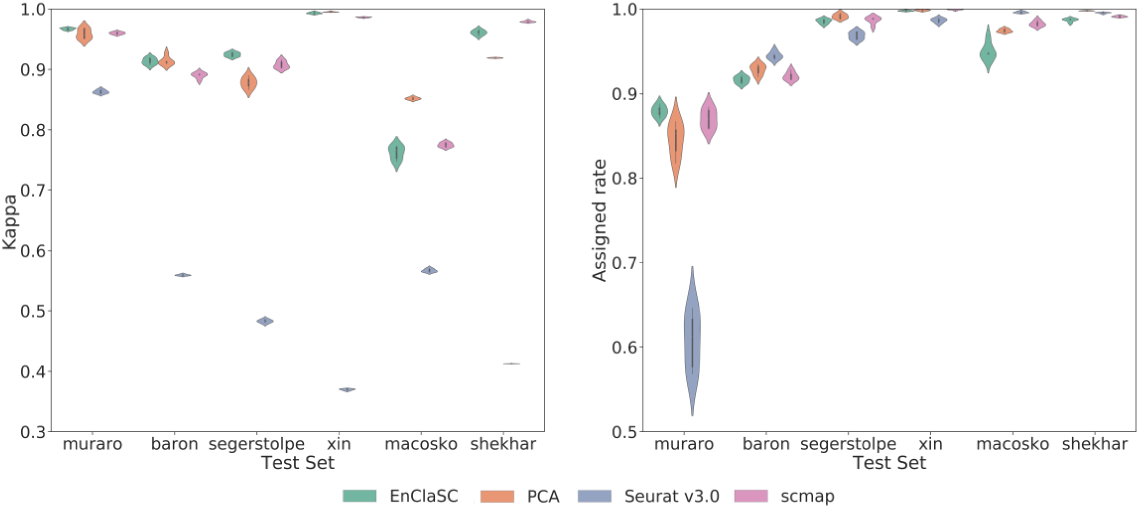
Performance of EnClaSC using different feature selection strategy.

#### Neural network module

We validated the contribution of the neural network module using the four human pancreatic datasets and the two mouse retina datasets, and we again performed six experiments as the above section. We replaced the neural networks in EnClaSC with a single neural network that does not use the ensemble learning strategy. We compared the performance of the above approach with that of our complete model, and show the results in Fig. 5. With 64.14% reduced variance of *kappa* and 71.97% reduced variance of *assigned rate* on average, the main contribution of ensemble learning strategy with several neural networks is significantly improving the stability of classification performance. Besides, on most datasets, the ensemble learning strategy slightly improves the classification performance.

**Fig. 5.**
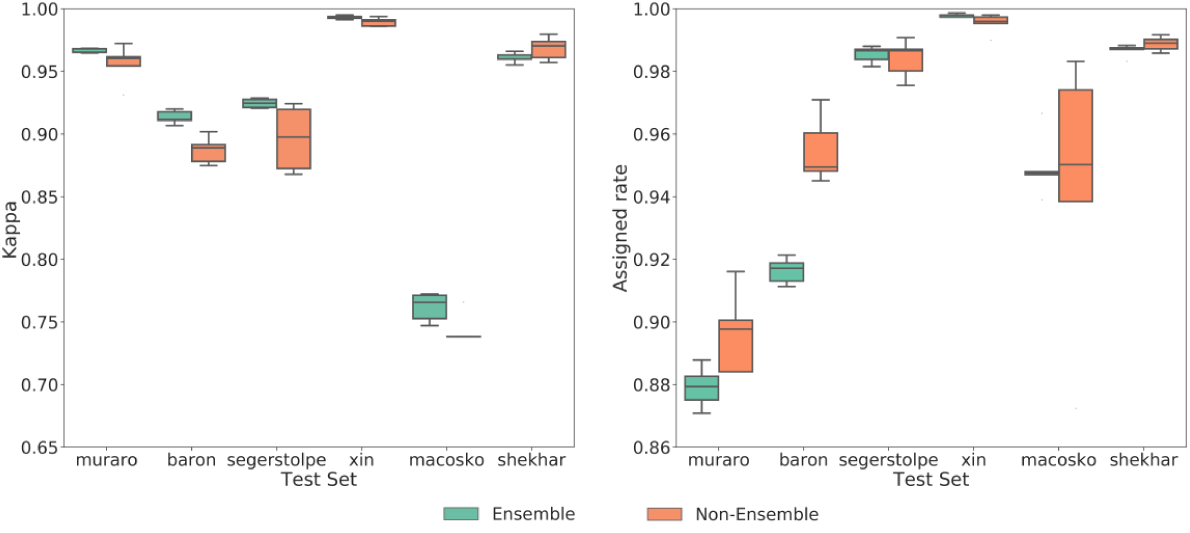
Performance of EnClaSC with or without ensemble learning in the neural network module.

#### Few-sample learning module

To demonstrate the contribution of the few-sample learning module, we compare the performance of EnClaSC with or without the few-sample learning module. We first used Shekhar Dataset as the training set, Macosko Dataset as the test set to evaluate the contribution of the few-sample learning module. This training-test set has a very distinct characteristic compared to other sets we have used, because there are massive samples of few-sample classes of the training set in the test set, which makes the classification task much more challenging. According to the definition in the Methods Section, we consider “rods” and “cones” as the few-sample classes in the training set. These two cell types account for 0.34% and 0.18% respectively in the training set, and account for 65.61% and 4.17% respectively in the test set.

As shown in Fig. 6a and 6b, EnClaSC with the few-sample learning module can identify 66.34% samples in the test set with the classification accuracy rate of 95.92%. However, the classification accuracy rate of EnClaSC without the few-sample learning module is only 87.33%. As shown in Fig. 6a and 6b, the number of incorrectly classified samples of the “rods” class is less than without the few-sample classification model, and the samples of the “cones” class in the test set are all classified incorrectly.

**Fig. 6.**
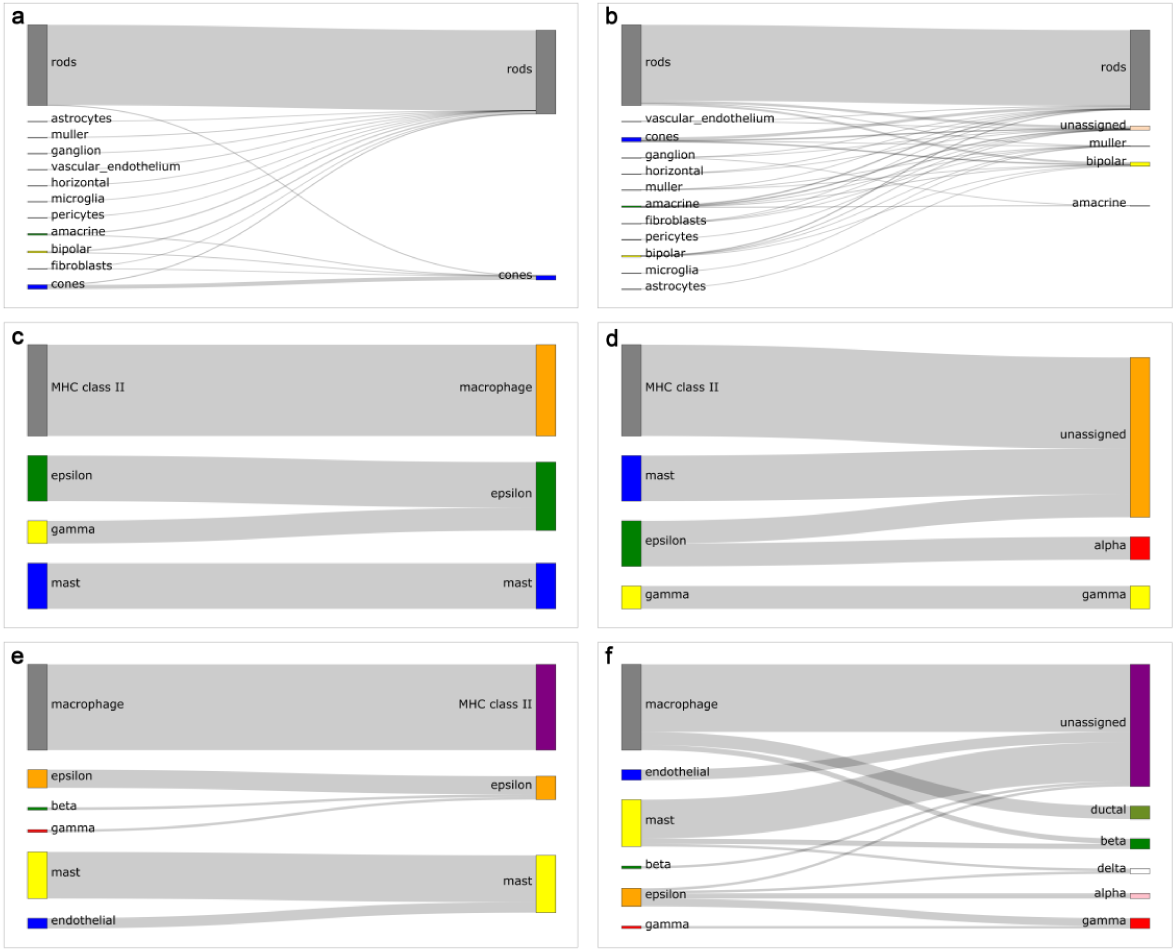
Performance of EnClaSC with or without the few-sample learning module. **a** The results of EnClaSC with few-sample learning module on Macosko Dataset. **b** The results of EnClaSC without few-sample learning module on Macosko Dataset. **c** The results of EnClaSC with few-sample learning module on Segerstolpe Dataset. **d** The results of EnClaSC without few-sample learning module on Segerstolpe Dataset. **e** The results of EnClaSC with few-sample learning module on Baron Dataset. **f** The results of EnClaSC without few-sample learning module on Baron Dataset.

We also evaluated the contribution of the few-sample learning module using training and test sets whose cell types have similar sample distributions. We integrated Baron Dataset, Muraro Dataset, and Xin Dataset as the training set, and used Segerstolpe Dataset as the test set, and Segerstolpe Dataset, Muraro Dataset and Xin Dataset as training sets, Baron Dataset as the test set to form two training-test sets for the experiment.

As shown in Fig. 6a and 6b, the training of few-sample classes in the neural network is insufficient with limited samples, resulting in the worse classification performance, while the few-sample learning module can effectively characterize the few-sample classes, and thus improve the classification performance.

In addition, we found that in the Segerstolpe Dataset, which contains “MHC class II” cell type but not ‘macrophage’ cell type. In the case that Baron Dataset or Segerstolpe Dataset serves as the test set, the few-sample learning module can associate the ‘macrophage’ class with the “MHC class II” class as shown in Fig. 6c-f. The data showed that “MHC class II” is a protein that is mainly secreted by the macrophage cell, indicating that the few-sample learning module possesses the potential ability to recognize new cell types and assign them to relative cell types. In summary, the few-sample learning module endows EnClaSC the ability to effectively resolve few-sample classes of the training set, while providing the potential ability to correlate new cell types with known relative cell types.

### EnClaSC scales up well with various data dimensionality

To illustrate the robustness of our method for various data dimensionality, we selected different numbers of features among the four human pancreatic datasets including Baron Dataset, Muraro Dataset, Xin Dataset, and Segerstolpe Dataset. We took each one of the four human pancreas datasets as a test set and integrated the remaining three datasets as a training set, and repeated each experiment five times. In order to illustrate the superior data-dimension adaptability of our method, we selected 100, 300, 500, 700 and 900 as the feature numbers of our method and scmap. It is worth noting that instead of designing a feature selection module, SuperCT directly binarizes all features, and thus uses all features in these experiments.

As shown in Fig. 7, the performance of EnClaSC slightly improves with the increase of data dimensionality. When the data dimensionality is 300, the *kappa* value of scmap rises to the peak and remains basically unchanged, while the *assigned rates* of scmap drops dramatically with the increase of data dimensionality. At the same time, we can see that when the data dimensionality reaches 500 or becomes higher, the *kappa* values and the *assigned rate* of EnClaSC are consistently higher than that of scmap and SuperCT. In summary, EnClaSC scales up well with various data dimensionality, and the higher the data dimensionality, the better the performance of EnClaSC.

**Fig. 7.**
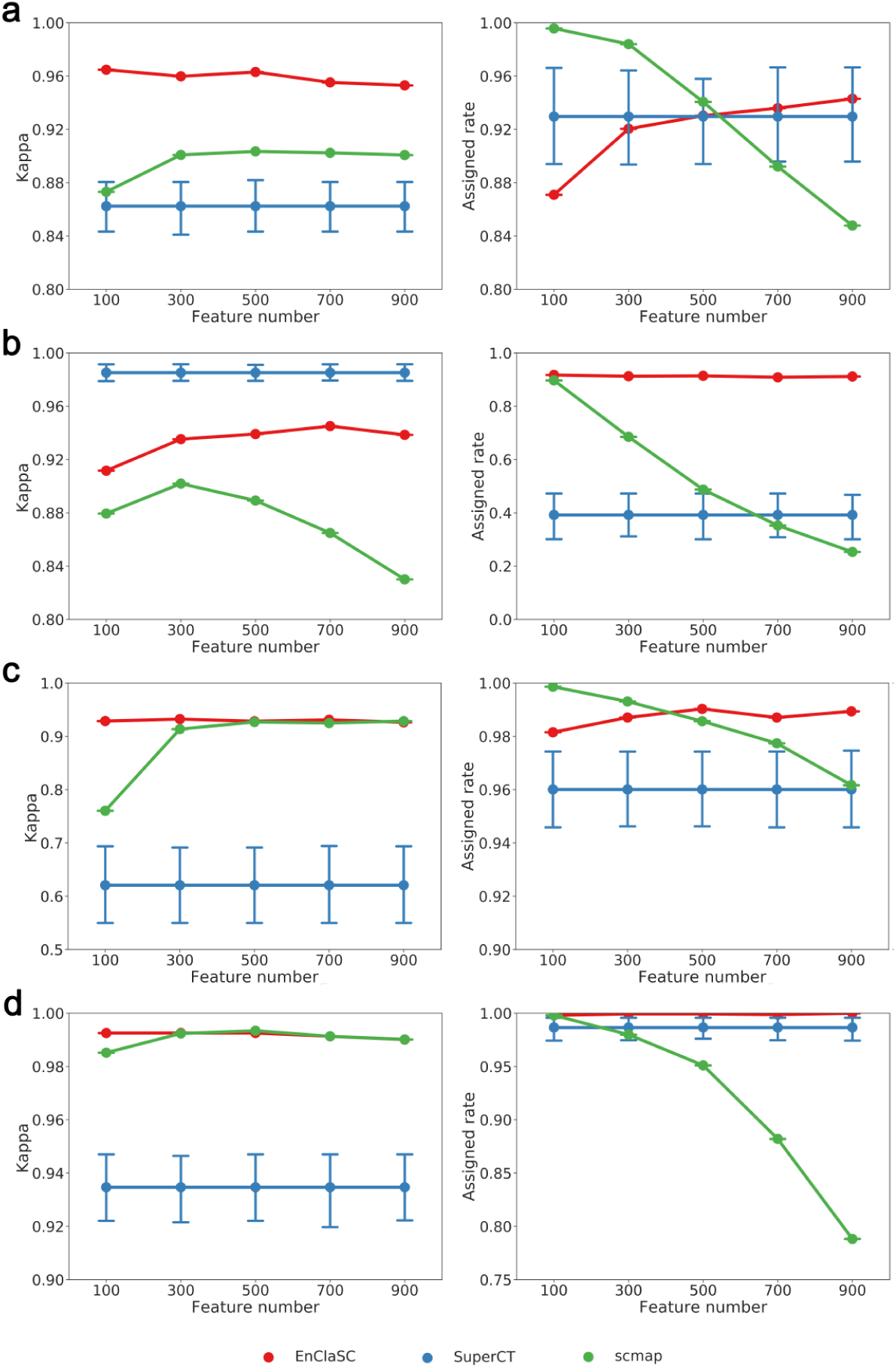
Performance comparison on datasets of various dimensionality. **a-d** The performance on Muraro Dataset, Baron Dataset, Segerstolpe Dataset and Xin Dataset, respectively.

### EnClaSC scales up well with different data sparsity

Considering that scRNA-seq data suffer from the dropout phenomenon, we further demonstrated that our method can scale up well with different data sparsity using the four human pancreatic datasets to perform four experiments as the above section. We randomly set 0%, 10%, 20%, 30%, 40%, and 50% of the non-zero elements in the raw expression matrix, and repeated each experiment five times. As illustrated in Fig. 8, EnClaSC is much more robust to the dropout rate compared with other two methods. The *kappa* values hardly change with the increasing of the dropout rate, even though the *assigned rate* decrease slightly. Nevertheless, the performance of scmap becomes worse because the randomly zeroing of the feature matrix could cause significant changes in cell-to-cell similarity. The performance of SuperCT fluctuates irregularly with the increasing dropout rate. In some cases, the *kappa* value may close to 0, indicating that the increasing dropout rate could cause further deterioration of the stability of SuperCT. The results demonstrate that EnClaSC can account for the sparsity of single-cell gene expression data, and thus benefits the cell-type classification tasks with the exponential growth of the Drop-seq based scRNA-seq data.

**Fig. 8.**
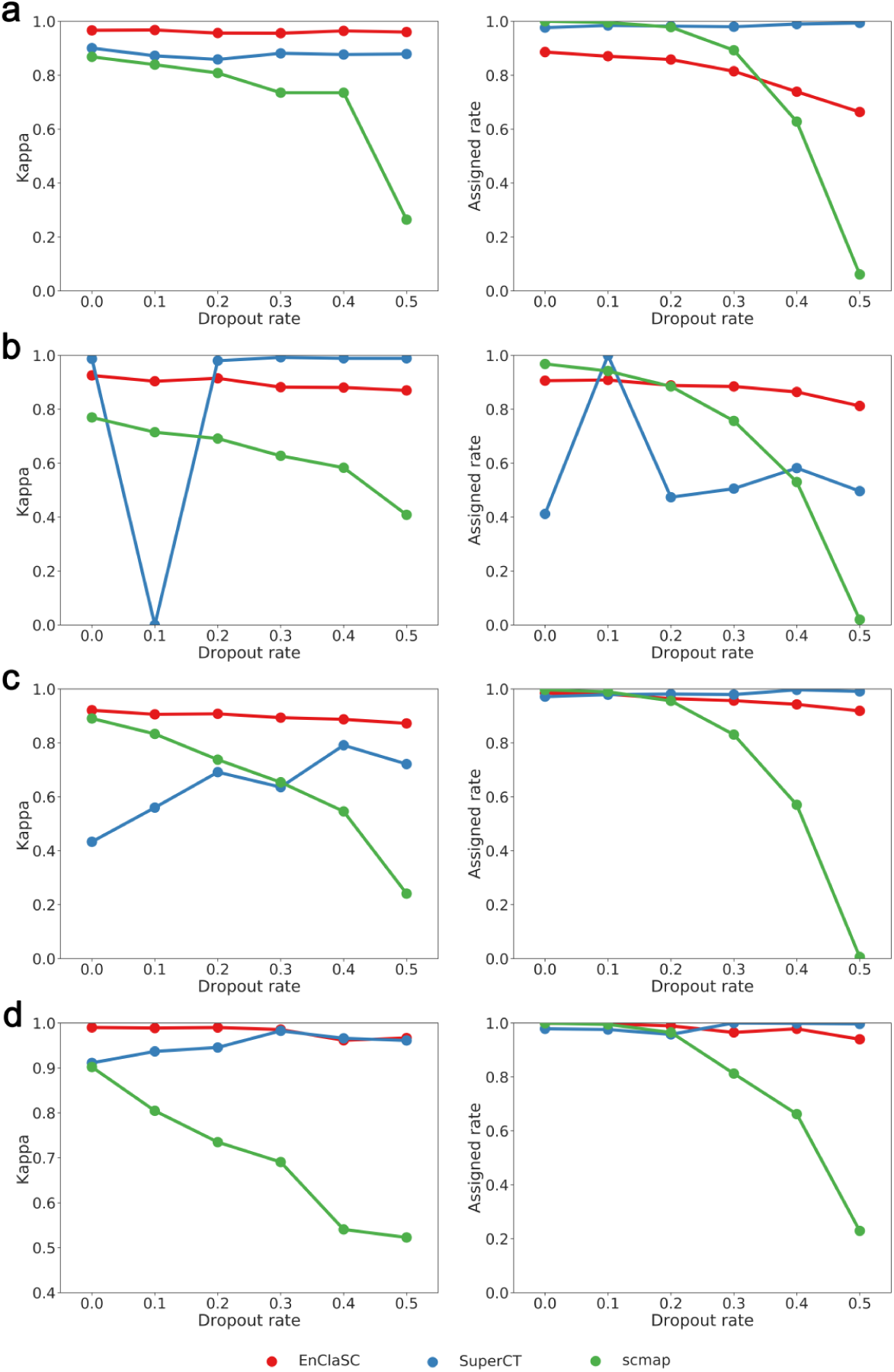
Performance comparison on datasets of different sparsity. **a-d** The performance on Muraro Dataset, Baron Dataset, Segerstolpe Dataset and Xin Dataset, respectively.

### EnClaSC enables cross-species classification

To illustrate that EnClaSC can effectively make **cross-species** classification, we used datasets of mouse brain cells provided by Romanov, R. A. et al. (Romanov Dataset for short) and Zeisel, A. et al. (Zeisel Dataset for short) as the training set, and a dataset of human brain cells provided by Darmanis, S. et al. (Darmanis Dataset for short) as the test set for the cross-species classification. Because there are only 466 samples on Darmanis Dataset, we did not use the Darmanis Dataset as the training set. As shown in Fig. 9, our method not only achieves the highest *assigned rate* (37.83% higher than scmap, and 2.54% higher than SuperCT), but also has the highest *kappa* (79.13% higher than scmap and 80.06% higher than SuperCT) in this cross-species classification task. Similarly, using our feature selection module in EnClaSC also provides the highest *kappa* value (211.60% higher than PCA, 11.89% higher than Seurat v3.0, and 4.94% higher than scmap), and *assigned rates* (0.51% higher than PCA, 2.84% higher than Seurat v3.0, and 2.46% higher than scmap). The results demonstrate that EnClaSC has superior classification performance both in the classification of homologous cells of the same species and in the classification of homologous cells of different species.

**Fig. 9.**
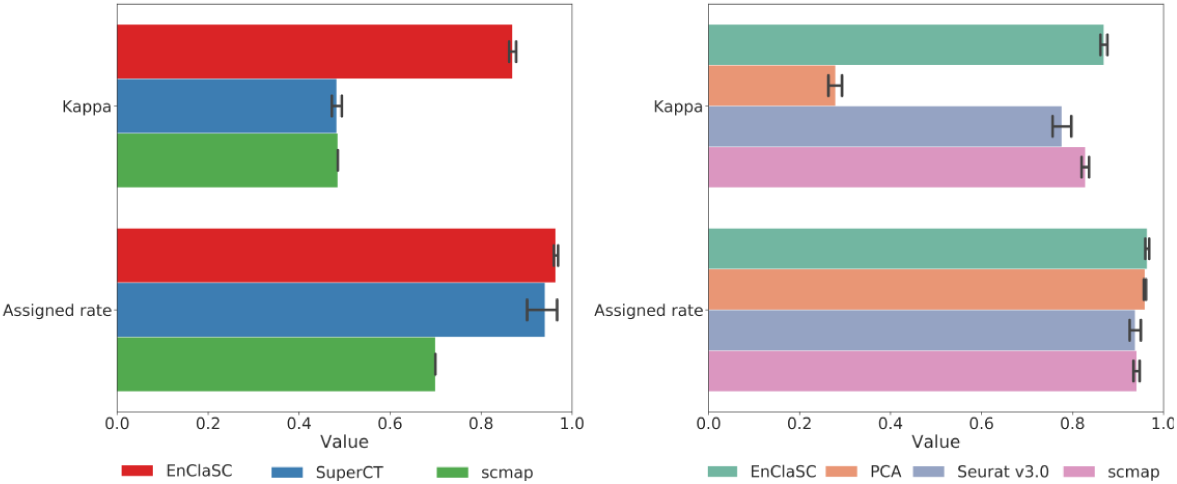
Performance comparison of cross-species classification.

## Discussion

scRNA-Seq techniques have advanced rapidly in recent years and enable the quantitative characterization of cell types at a single-cell resolution. EnClaSC is proposed to classify cell types using supervised learning. With the contribution of the tailored feature selection, neural network and few-sample learning modules, our method is superior to other baseline methods, such as scmap and SuperCT, with regard to not only accuracy and robustness, but also the performance of classifying few-sample classes.

Our method can certainly be improved in some aspects. First, a more well-crafted module, such as modules considering dropout evens, can be introduced to better characterize the scRNA-seq data, and thus further improves the performance of our method. Second, our method can be extended to incorporate other types of functional genomics data such as chromatin accessibility [23, 24]. Finally, drawing on the idea of VPAC, we can integrate the feature selection module with other modules to endow the method with the ability to balance the feature selection and prediction steps, and thus extract features that are more conducive to the cell type classification [25].

## Conclusions

We have proposed a supervised learning method, named EnClaSC, for accurate and robust cell-type classification of single-cell transcriptomes. Each of the well-crafted feature selection, neural network and few-sample learning modules draws on the idea of ensemble learning, which makes EnClaSC superior to existing methods in the self-projection within a specific scRNA-seq dataset, the cell-type classification across different scRNA-seq datasets, various data dimensionality, and different data sparsity. We have further demonstrated the ability of EnClaSC to effectively make cross-species classification, which may shed light on the studies in the correlation of different species. Eventually, we expect that such a supervised learning approach will be widely applicable for the cell-type classification with the explosive growth of scRNA-seq data.

## Abbreviations

scRNA-seq: single-cell RNA-sequencing
ANN: artificial neural network
PCA: principal component analysis

## Declarations

### Ethics approval and consent to participate

Not applicable.

### Consent for publication

Not applicable.

### Availability of data and material

The datasets supporting the conclusions of this article are publicly available from the NCBI Gene Expression Omnibus (https://www.ncbi.nlm.nih.gov/geo/), the NCBI Sequence Read Archive (http://www.ncbi.nlm.nih.gov/Traces/sra/), and the EMBL-EBI ArrayExpress (https://www.ebi.ac.uk/arrayexpress/).

### Competing interests

The authors declare that they have no competing interests.

### Funding

This research was partially supported by the National Key Research and Devel-opment Program of China (No. 2018YFC0910404), the National Natural Science Foundation of China (Nos. 61873141, 61721003, 61573207), and the Tsinghua-Fuzhou Institute for Data Technology.

### Authors’ contributions

RJ designed the research. XC and SC designed and implemented the models. XC collected data and analyzed the results. XC, SC and RJ wrote the manuscript. All authors read and confirmed the manuscript.

## Acknowledgements

Rui Jiang is a RONG professor at the Institute for Data Science, Tsinghua University.

